# Effects of external stimulation on psychedelic state neurodynamics

**DOI:** 10.1101/2020.11.01.356071

**Authors:** Pedro A.M. Mediano, Fernando E. Rosas, Christopher Timmermann, Leor Roseman, David J. Nutt, Amanda Feilding, Mendel Kaelen, Morten L. Kringelbach, Adam B. Barrett, Anil K. Seth, Suresh Muthukumaraswamy, Daniel Bor, Robin L. Carhart-Harris

**Affiliations:** Department of Psychology, University of Cambridge, Cambridge CB2 3EB; Psychedelic Research Center, Department of Brain Science, Imperial College London, W12 0NN, UK; Data Science Institute, Imperial College London, London SW7 2RH; Center for Complexity Science, Imperial College London, London SW7 2RH; The Beckley Foundation, Beckley Park, OX3 9SY, Oxford, United Kingdom; Wavepaths, London E1 5JL, UK; Department of Psychiatry, University of Oxford, Oxford OX3 7JX, UK; Center for Music in the Brain, Department of Clinical Medicine, Aarhus University, DK-8000 Aarhus C, Denmark; Sackler Center for Consciousness Science and Department of Informatics, University of Sussex, Brighton BN1 9RH, UK; CIFAR Program on Brain, Mind, and Consciousness, Toronto M5G 1M1, Canada; School of Pharmacy and Centre for Brain Research, Faculty of Medical and Health Sciences, The University of Auckland, Auckland, New Zealand

**Keywords:** Complexity, Psychedelics, Neuroscience, Consciousness

## Abstract

Recent findings have shown that psychedelics reliably enhance brain entropy (understood as neural signal diversity), and this effect has been associated with both acute and long-term psychological outcomes such as personality changes. These findings are particularly intriguing given that a decrease of brain entropy is a robust indicator of loss of consciousness (e.g. from wakefulness to sleep). However, little is known about how context impacts the entropy-enhancing effect of psychedelics, which carries important implications for how it can be exploited in, for example, psychedelic psychotherapy. This article investigates how brain entropy is modulated by stimulus manipulation during a psychedelic experience, by studying participants under the effects of LSD or placebo, either with gross state changes (eyes closed vs. open) or different stimulus (no stimulus vs. music vs. video). Results show that while brain entropy increases with LSD in all the experimental conditions, it exhibits largest changes when subjects have their eyes closed. Furthermore, brain entropy changes are consistently associated with subjective ratings of the psychedelic experience, but this relationship is disrupted when participants are viewing video — potentially due to a “competition” between external stimuli and endogenous LSD-induced imagery. Taken together, our findings provide strong quantitative evidence for the role of context in modulating neural dynamics during a psychedelic experience, underlining the importance of performing psychedelic psychotherapy in a suitable environment. Additionally, our findings put into question simplistic interpretations of brain entropy as a direct neural correlate of conscious level.

**Significance Statement:** The effects of psychedelic substances on conscious experience can be substantially affected by contextual factors, which play a critical role in the outcomes of psychedelic therapy. This study shows how context can modulate not only psychological, but also neurophysiological phenomena during a psychedelic experience. Our findings reveal distinctive effects of having eyes closed after taking LSD, including a more pronounced change on the neural dynamics, and a closer correspondence between brain activity and subjective ratings. Furthermore, our results suggest a competition between external stimuli and internal psychedelic-induced imagery, which supports the practice of carrying out psychedelic therapy with patients having their eyes closed.

---

**P**sychedelic substances, such as LSD and psilocybin, are known to induce profound changes in subjects’ perception, cognition, and conscious experience. In addition to their role in ancestral spiritual and religious practices, and their recreational use related to introspection and self-exploration, there is promising evidence that psychedelics can be used therapeutically to treat multiple mental health conditions (1–4). However, despite the increasingly available evidence of the neurochemical action of psychedelics at the neuronal and sub-neuronal level (5, 6), the mechanisms associated with their therapeutic efficacy are not yet completely understood.

Some of the factors at play during psychedelic therapy can be related to the Entropic Brain Hypothesis (EBH) (7, 8), a simple yet powerful theory which posits that the rich altered state of consciousness experienced under psychedelics depends on a parallel enriching effect on the dynamics of spontaneous population-level neuronal activity.^*^ The hypothesis that increased brain entropy — as captured e.g. by Lempel-Ziv (LZ) complexity (8) — corresponds to states of enriched experience has found empirical support in neuroimaging research on psychedelics (9, 10), as well as on other altered states, like meditation (11) and states of “flow” associated with musical improvisation (12). Furthermore, the therapeutic mechanisms of psychedelics are thought to depend on their acute entropy-enhancing effect, potentially reflecting a window of opportunity (and plasticity) mediating therapeutic change (13, 14). Conversely, states such as deep sleep, general anaesthesia, and loss of consciousness have consistently shown reduced brain 28 entropy (15–17).

The effectiveness of psychedelic therapy is thought to depend not only on direct neuropharmacological action, but also on contextual factors — commonly referred to as *set and setting*. These include the subject’s mood, expectations, and broader psychological condition (set) prior to the “trip”, together with the sensorial, social, and cultural environment (setting) in which the drug is taken. For example, there is direct physiological evidence that (visual) stimuli affect the expression of serotonergic receptor genes (18), and that specific music choices may either enhance or impede therapeutic outcomes (19).

Despite its presumed importance, to our knowledge no previous study has systematically assessed the influence of set and setting on brain activity and subjective experience during a psychedelic experience. This lack of relevant research, combined with the fact that psychedelic therapy is almost exclusively carried out with music listening and eyes closed, exposes a knowledge gap that compromises key assumptions of current psychedelic therapy practice. Here, we provide a first step towards bridging this gap, presenting a systematic investigation of how different environmental conditions can modulate changes in brain entropy elicited by psychedelics in healthy subjects. This work provides a proof of principle that paves the way for future studies with clinical cohorts.

## Results

### Increased LZ under external stimulation

We use data presented by Carhart-Harris *et al*. (20), together with previously unpublished data from the same experiment. Twenty subjects participated in the study by attending two experimental sessions: one in which they received intravenous (i.v.) saline (placebo), and one in which they received i.v. LSD (75μg). The order of the sessions was randomised, separated by two weeks, and participants were blind to the order (i.e. single blind design). Whole-brain magnetoencephalography (MEG) data were collected under four conditions: resting state with eyes closed, listening to instrumental ambient music with eyes closed, resting state with eyes open (focusing on a “fixation dot”), and watching a silent nature documentary video — henceforth referred to as *closed, music, open*, and *video*. The music tracks were taken from the album “Eleusian Lullaby” by Alio Die, and the video was composed of segments of the “Frozen Planet” documentary series produced by the BBC. More information about the experimental design can be found in Ref. (20).

Studying the whole-brain average LZ from the placebo sessions showed that external stimuli yield significant differences in LZ (Kruskal-Wallis test, *p*< 0.001). Post-hoc t-tests, shown in Figure 1a, revealed that richer stimuli induce consistent significant increases across conditions, with large effect sizes (Cohen’s *d*).

**Fig. 1.**
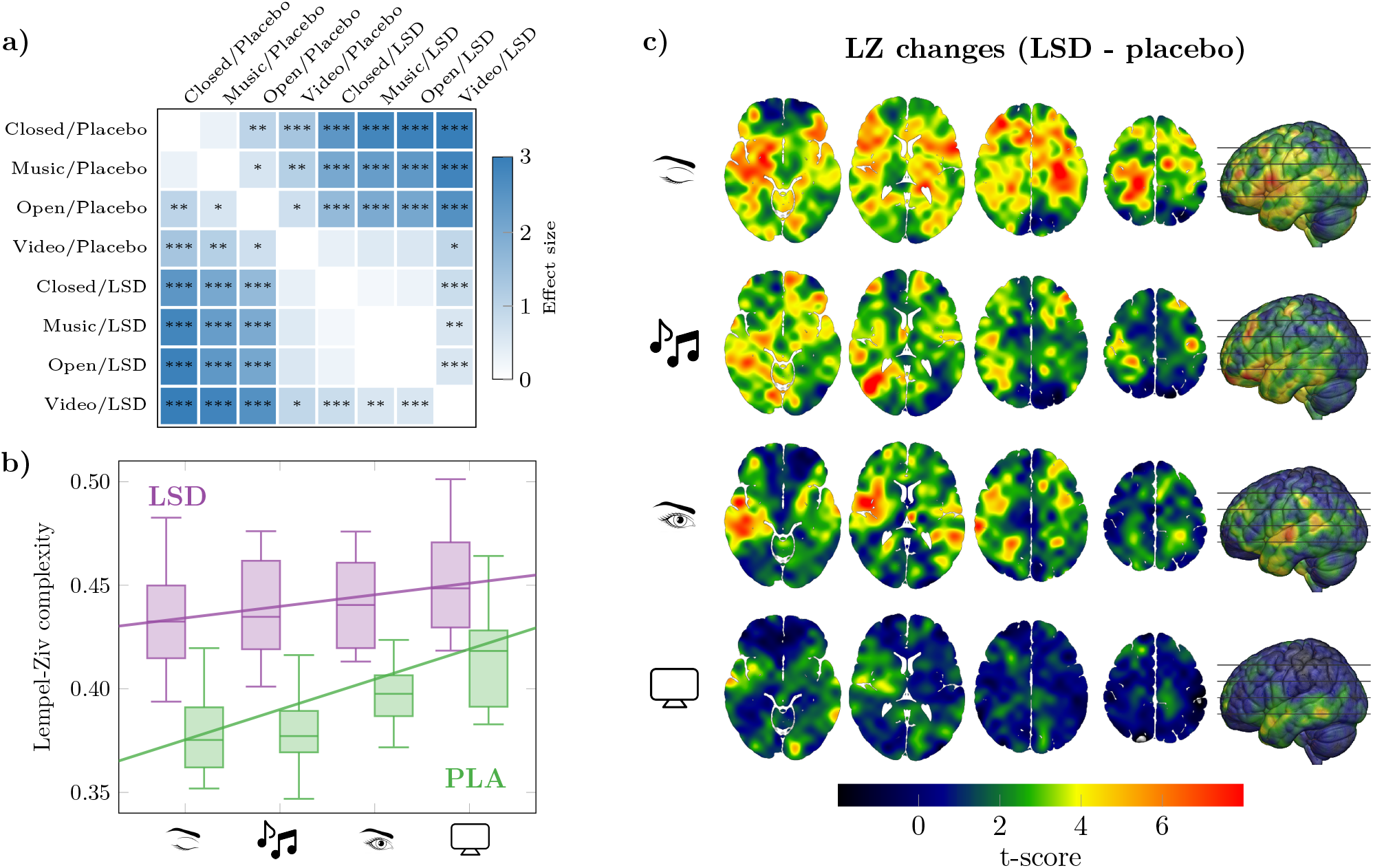
Stronger external stimulation increases baseline entropy, reduces drug effect. **a)** The differences in average LZ, as measured by post-hoc t-tests and effect sizes (Cohen’s d), increase with stimulus and drug (*: *p* < 0.05,**: *p* < 0.01, ***: *p* < 0.001). **b)** However, stronger external stimulation (i.e. with higher baseline LZ) *reduces* the differential effect of LSD on brain entropy vs. placebo. Linear mixed-effects models fitted with LZ complexity as outcome show a significant negative drug × condition interaction (*p* < 0.01; see Supplementary Material). **c)** T-scores for the effect of the drug in all four experimental conditions. In agrement with the LME models, the effect of the drug on increasing LZ substantially diminishes with eyes open or under external stimuli.

To disentangle the effect of the stimuli over the effect of eye opening, a linear mixed-effects (LME) model was constructed using the presence of stimulus and eye opening as predictor variables, and subject identity as random effect (see Methods). This model showed significant positive effects of both stimulus (*β* = 0.013, SE = 0.005, *p* = 0.017) and eye opening (*β* = 0.025, SE = 0.005, *p* < 0.001). The statistical significance of both effects suggests that the measured LZ cannot be explained merely by the presence or absence of visual stimuli, and must be related to the structure of such stimuli (either music or the video). Nonetheless, it is noteworthy that the simple act of opening one’s eyes has an especially marked (augmenting) effect on brain entropy.

### Stronger external stimulus weakens drug effect

To study the effect of LSD on the whole-brain average LZ, we constructed LME models similar to those in the previous section and added the drug as fixed effect. This analysis shows a dramatic increase in LZ under the effects of LSD, much larger than that associated with eye opening or stimulus (Fig. 1b and Table 1). Post-hoc analyses showed that the effect of drug is substantial in all stimulus conditions (Fig. 1a).

**Table 1.** Means (*β*) and standard errors (SE) of coefficients of the LME model predicting whole-brain average LZ.

| | $\beta$ | SE | $p$ |
| --- | --- | --- | --- |
| **Drug** | 0.047 | 0.005 | <0.001 |
| **Stimulus** | 0.010 | 0.003 | 0.002 |
| **Eyes** | 0.025 | 0.005 | <0.001 |
| **Drug × Eyes** | -0.016 | 0.006 | 0.011 |

Crucially, the LME model revealed a significant interaction between drug and eye opening as predictors of LZ (Table 1). Importantly, this interaction effect was negative — i.e. increased external stimulation *reduced* the effect of the drug. Alternatively, this can be interpreted as the drug reducing the effect of external stimulation on brain entropy — which, either way, points towards a “competition” between endogenous, drug-induced, and exogenous, stimulus-induced, effects on neural dynamics (21). This negative interaction was confirmed by ordering the four experimental conditions with integer values from 1 to 4 (Fig. 1b), and with multiple statistical hypothesis tests (e.g. 2-way ANOVA). Furthermore, we confirmed that the results still hold with stricter filters (e.g. a low-pass filter at 30 Hz on the MEG signals), and when controlling for order effects between the stimulus and non-stimulus sessions (see Supplementary Material). Both the effect of the drug and its interaction with external conditions are spatially widespread (Fig. 1c).

### Setting modulates subjective ratings and their relationships

In addition to MEG measurements, Visual Analog Scale (VAS) subjective ratings were collected at the end of each session. The questionnaires were designed to capture central features of the subjective effects of LSD. They included assessments of the intensity of the experience, emotional arousal, ego dissolution, positive mood, and simple and complex internal visual imagery. The imagery items were only rated for the eyes-closed conditions.

The effects of LSD on VAS ratings varied widely between conditions (Fig. 2a). A quantitative analysis with LME models showed the effect of the drug to be much larger than that of the stimulus or eye opening on all the VAS measures (Fig. 2b). Additionally, stimulus effects tended to be more specific than drug effects, reaching statistical significance only for positive mood and emotional arousal — in line with previous findings that carefully selected stimuli (e.g. music) can boost the affective state of subjects undergoing psychedelic psychotherapy (19, 22, 23). It is worth noting that these two are the least psychedelic-specific items.

**Fig. 2.**
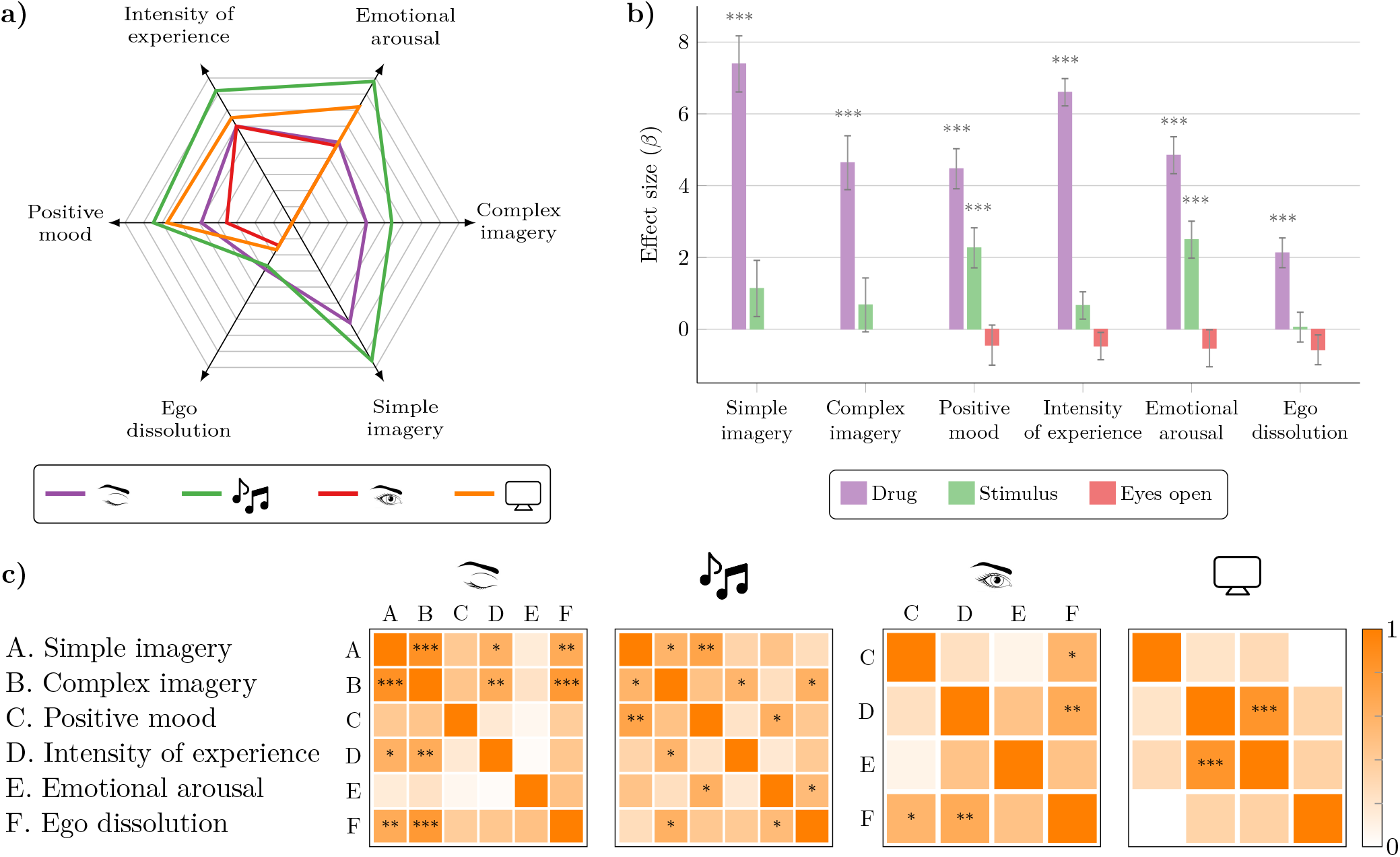
Setting affects participants’ subjective reports of their psychedelic experience. **a**) Average increases in VAS ratings between LSD and placebo show a varied profile across experimental conditions, suggesting that setting modulates participants’ rating of their own experience. Simple and complex imagery data was not collected in the eyes open and video conditions. **b**) Effect sizes obtained from LME modelling confirm a strong effect from the drug in all items, as well as smaller and more specific effects from stimulus. **c**) Between-subjects correlation matrices between experience reports. (*: *p* < 0.05,**: *p* < 0.01,***: *p* < 0.001)

Differences in setting not only affected subjects’ VAS ratings, but also the relationship between the ratings themselves (Fig. 2c). For example, when resting with eyes closed, subjects tended to rate the intensity of their experience in agreement with the vividness of their simple and complex imagery — but, when watching a video, intensity was more strongly correlated with emotional arousal. These findings show that what subjects consider their intensity of experience can dramatically vary across various dimensions (24), confirming the assumption that the subjective quality and general intensity of a psychedelic experience strongly depends on the environmental conditions (or setting) in which it takes place.

### Neural-psychometric correlations can be disrupted by external stimuli

A major aim of psychedelic neuroimaging is to discover specific relationships between brain activity and subjective experience. Examples include mappings between specific neural dynamics and ratings of ego dissolution (25) or other specific aspects of experience such as its visual quality (9). However, given that — as we show here — setting interacts with neural dynamics, then it is natural to ask whether it also affects the relationship between phenomenology and its neural correlates.

To address this question, we analysed the relationship between LZ and VAS changes induced by LSD, in each one of the four experimental conditions. Between-subjects Pearson correlation coefficients were calculated between changes in VAS ratings and LZ measured in different regions of interest (ROI). Motivated by the nature of the study and known brain effects of LSD (20, 25), we focused on areas associated with sensory processing (visual and auditory), interoception (insula), emotional processing (amygdala), and self-monitoring (mPFC and posterior DMN; see Methods for details).

Analyses revealed multiple significant relationships between subjective ratings and LZ changes during the eyes-closed, music, and eyes-open conditions (Fig. 3). For example, we observed significant (*p* < 0.05, FDR-corrected) positive correlations between ego dissolution and DMN, positive mood and amygdala, and simple and complex imagery and visual and auditory ROIs, all in the eyes-closed condition — supporting the suitability of the eyes-closed resting condition for assessing the neural correlates of these experiences. Strikingly, all the observed neural-psychometric correlations vanish when subjects watched a video, with none exceeding an absolute value of |*r*| > 1/10. This observation was verified by building a multivariate regression model, using the correlation coefficients between VAS and LZ changes as target variables, and stimuli and eye opening as predictors. Results showed that neither stimuli (*p* = 0.17) nor eyes-open (*p* = 0.13) had significant effects by themselves, but their interaction was strongly associated with smaller VAS-LZ correlation values (*β* = −0.21, SE = 0.08, *p* = 0.006; see Supplementary Material).

**Fig. 3.**
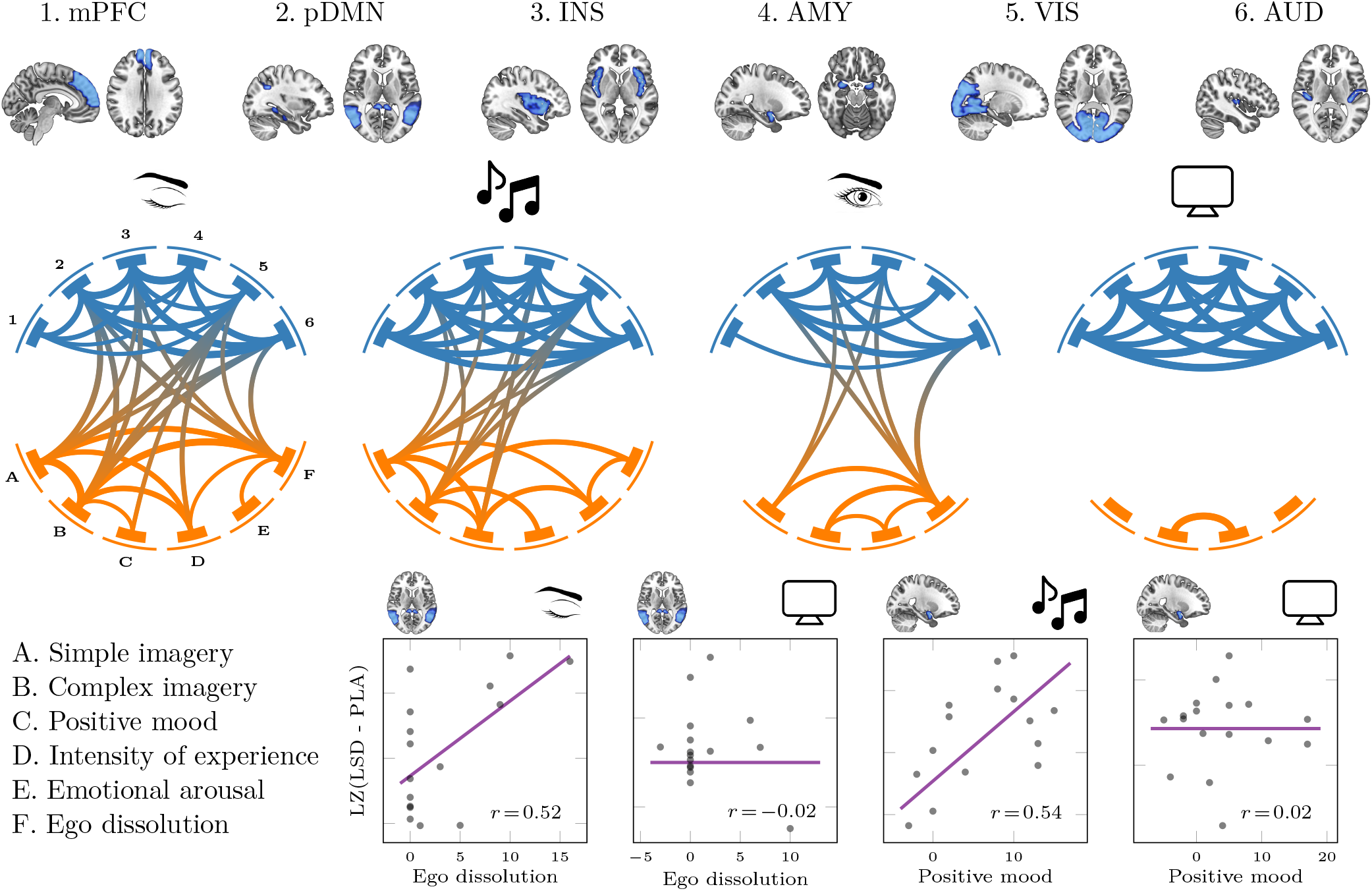
External stimulation alters the relationship between psychometric and neural effects of LSD. Network representation of correlation matrices between brain entropy in six regions of interest (numbered 1-6, top), and subjective experience ratings (labelled A-F, bottom left), in all four experimental conditions. As external stimulation is increased, there is a large decrease in correlation between subjective ratings and entropy, but an increase in the correlation in entropy between different brain regions (see Supplementary Material). Bottom right panels show example correlations between ego dissolution and posterior DMN entropy (two left panels) and positive mood and amygdala entropy (two right panels). In both cases, correlation is strong and significant with eyes closed, but vanishes when subjects watch a video.

As a complementary analysis, we also studied how the four environmental conditions affect the relationship between the LSD-induced LZ changes across different ROIs. To do this, we evaluated the Pearson correlation coefficient between the LZ changes measured in the various ROIs across subjects. It was observed that the correlation between ROIs is substantially increased when subjects perceive an external stimulus (either music or video; see Supplementary Material), which could be indicative of a form of “complexity matching” (26) in which neural dynamics are entrained by the external stimulus, obscuring the relationship between neurodynamics and subjective experience. This observation was also verified via multivariate regression modelling, this time using ROI-ROI correlation values as target. In this case, eye opening was associated with smaller correlation values (*β* = −0.10, SE = 0.04, *p* = 0.011), while stimuli (*β* = 0.15, SE = 0.04, *p* < 0.001) and the interaction between stimuli and eyes-open (*β* = 0.18, SE = 0.05, *p* = 0.001) were both associated with significantly larger correlation values (see Supplementary Material). These findings suggest that the increased within-brain correlation driven by external stimulation may obfuscate potential correlations between entropy and individual VAS ratings — which are most apparent e.g. in the eyes closed condition.

### Conditional predictive analyses of subjective reports

Finally, we analysed the relationship between changes in LZ and behavioural reports as they were exposed to the different experimental conditions. For this, we constructed LME models using VAS ratings as target; average LZ, eye opening, and stimulus as fixed effects; and subject identity as random effect (see Methods).

These models revealed multiple associations between brain entropy and subjective reports (Fig. 4), including some widespread correlations with LZ averaged across the whole brain (most strongly with ego dissolution and simple imagery), as well as more specific correlations (e.g. between positive mood and amygdala). In contrast, stimulus and eye opening show small effect sizes in all models, as well as strong negative interactions with LZ (see Supplementary Material). This negative interaction suggests that the relationship between LZ and VAS is broken when stimuli are present, in line with the results in Fig. 3.

**Fig. 4.**
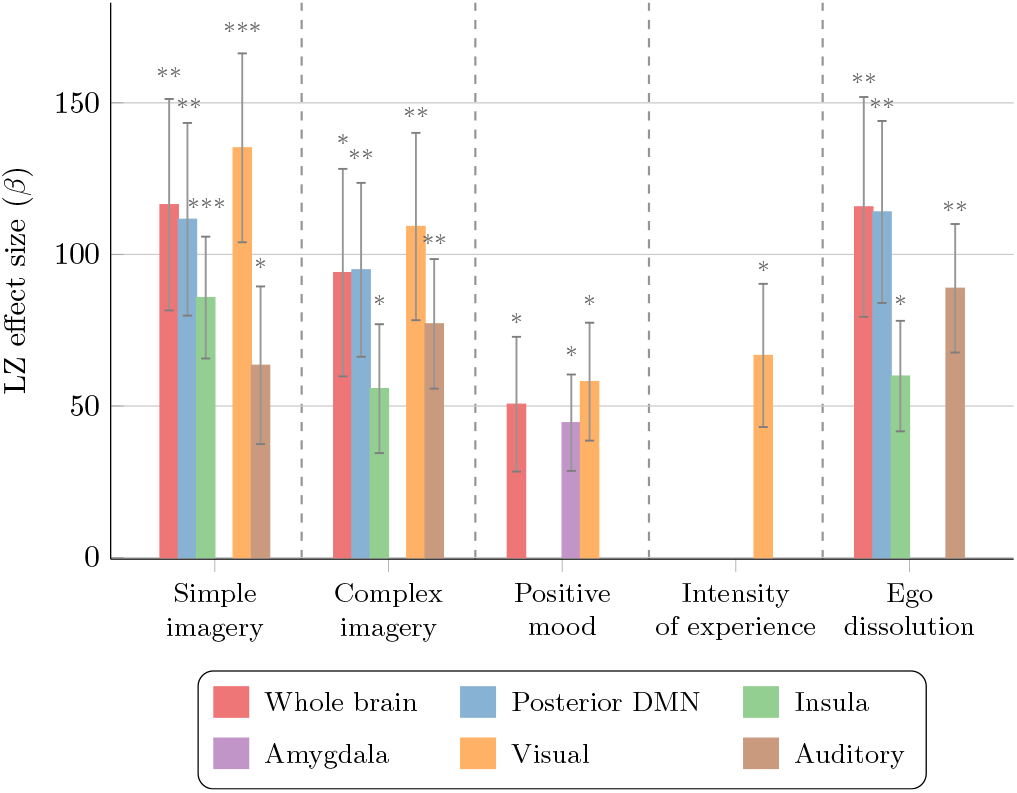
Changes in brain entropy predict changes in subjective reports. Estimates, standard error, and FDR-corrected statistical significance (*: *p* < 0.05,**: *p* < 0.01,***: *p* < 0.001), of the effect of the LZ differences (LSD-PLA) for predicting VAS differences (LSD-PLA), obtained from LME models calculated over the four conditions.

To explore the correlations between behavioral ratings and LZ in various ROIs in more detail, we performed a conditional predictive power analysis (see Methods). This method allows us to build a directed network representing the predictive ability of the various ROIs with respect to a given VAS item, such that a ROI *R*_1_ is connected to a VAS item *V* via a another ROI *R*_2_ if, once the entropy change in *R*_2_ is known, there is no further benefit in knowing the entropy change in *R*_1_ for improving the prediction of the change in *V* (Fig. 5a). Results show that, in general, “low-level” regions (i.e. closer to the sensory periphery, like visual areas) tend to “mediate” the associations between subjective reports and high-level regions (like the DMN). For example, visual and auditory areas mediate the predictive information that the pDMN and insula have about reported complex imagery.^†^ Put simply: once the change in entropy in auditory and visual regions is known, knowing the change in entropy in the pDMN provides no extra information about the change in reported complex imagery. A notable exception, however, is ego dissolution, for which pDMN, auditory, and insula all provide unmediated complementary information — in line with previous studies linking self-related processing and the DMN (20).

**Fig. 5.**
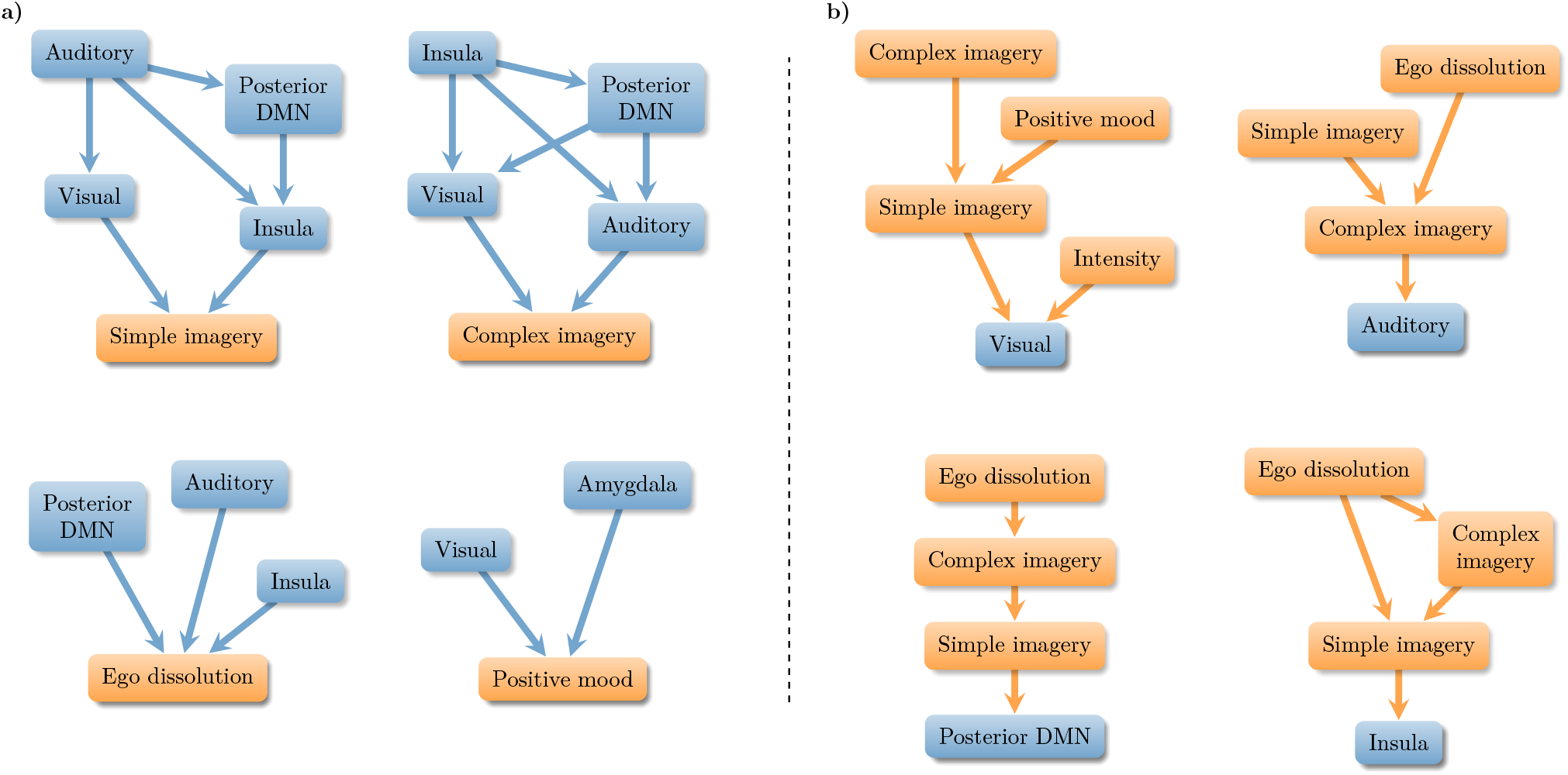
Statistical structure of brain entropy and subjective ratings data. Networks represent conditional prediction diagrams (see Methods), in which node *i* is connected to node *j* if *j* “mediates” the statistical predictive information that *i* has about a target variable (bottom node in each network). **a)** Conditional predictive analysis from brain entropy to subjective experience reports; and **b)** from subjective reports to brain entropy.

We also performed a reciprocal analysis to assess the conditional predictive power of the various VAS items using LZ as target (Fig. 5b). Results show that, across brain regions and VAS items, the predictive power of more abstract VAS scores (e.g. ego dissolution, positive mood) tends to be mediated by less abstract ones (e.g. simple and complex imagery). For example, changes in ego dissolution scores become irrelevant for predicting LZ in auditory areas once one knows the corresponding change in complex imagery. One interpretation of these analyses is that brain entropy, as currently measured with LZ, may most faithfully reflect “low-level” aspects of the brain-mind relation (see Discussion).

## Discussion

The present study’s findings provide strong quantitative evidence on how environmental conditions can have a substantial influence on both subjective experience and on neural dynamics during a psychedelic experience. Importantly, the entropy-enhancing effects of LSD were less marked when participants opened their eyes or perceived external stimuli — such as music or video. Furthermore, the differences in brain entropy observed in various regions of the brain were found to be associated with behavioural reports about the subjects’ perception, emotion, and self-related processing — but the relationship between brain entropy and subjective reports collapsed in the video-watching condition.

### LZ as a robust correlate of subjective experience

The increase in brain entropy — seen via LZ — is known to be a robust M/EEG biomarker associated with the psychedelic state (9, 10), and indeed, conscious states more generally (15–17, 27). In addition to replicating this effect on new data, we also observed other known effects of serotonergic psychedelics, including pronounced spectral power changes (in particular, LSD-induced alpha suppression (28)). Interestingly, the relationship between changes in these other metrics (like alpha power) and subjective ratings was substantially weaker than that of LZ (see Supplementary Material), suggesting that LZ is a particularly well-suited marker of psychedelic subjective experience.^‡^

Our results further put into question interpretations of LZ as a simple correlate of overall conscious level (e.g. (30)). In effect, subjects under LSD watching a video had the highest absolute brain entropy, but did not give maximal subjective ratings in any of the psychometric items. Furthermore, while a profound subjective experience such as ego dissolution was found to correlate with LZ changes, this effect was found most prominently in the eyes closed condition, and its predictive power was mediated by reported (simple and complex) visual imagery.

We propose two alternative interpretations of these findings. On the one hand, it could be that LZ most faithfully indexes brain activity associated with low-level sensory processing. This interpretation could be seen as consistent with recent reports showing that LZ is not affected by cognitive load (31). On the other hand, it could be that LZ shows strong associations with high-level cognitive processing or subjective phenomena (such as ego dissolution) in the eyes-closed conditions because that relationship becomes more specific in the absence of the strong “driving” effects present in the eyes-open conditions — especially video. Future studies might distinguish between these hypotheses by exploring the reliability of relationships between LZ and various subjective phenomena, including ego dissolution, perceptual complexity, and alertness, involving different pharmacological agents (e.g. psychedelics and stimulants), dosages, and stimuli.

### Towards a refinement of the entropic brain hypothesis

A deeper understanding of the functional relevance of brain entropy will help us better understand how such measures can be refined, in order to shed clearer light on their relationship with reported phenomenology. The results presented in this paper, while grounded in and motivated by the EBH, also highlight some important qualifiers of it. Since brain entropy measures such as LZ depend only on the dynamics of individual loci (e.g., individual time series corresponding to single sources or sensors), they may only indirectly reflect the richer scope of brain dynamics, network and connectivity properties — although it is worth noting that LSD-induced entropy increases at the single-source level have been related to specific network properties of the human connectome (32).

One potential way forward for the EBH may be to consider the entropy of network dynamics and other high-order brain features, rather than merely the entropy of individual sources. For example, examining increases in entropy at the level of emergent whole-brain states may prove particularly fruitful (33). We see this as part of a broader move towards multidimensional descriptions of brain activity, transcending “one-size-fits-all” scalar measures — including more complicated unidimensional ones like integrated information (34, 35). In line with recent theoretical proposals (36) and experimental findings (10), a range of metrics may be necessary to provide a more complete, multi-dimensional representation of brain states. However, we also acknowledge that increasing model complexity can complicate interpretability and affect statistical power, and thus is only justified when it yields substantial improvement in explanatory power and is driven by reliable hypotheses.

### Implications for psychedelic psychotherapy

These findings can be regarded as neurobiological evidence for the importance of environmental context (37), or ‘setting,’ to the quality of psychedelic experiences — a matter of particular relevance to psychedelic therapy. In particular, the present findings support the principle that having one’s eyes closed during a psychedelic experience may enhance the differential entropic effect of the drug (1), which is consistent with approaches fostering eyes-closed, introspective experiences during psychedelic therapy, as they may lead to beneficial therapeutic outcomes (38). In addition, our results suggest a differential effect between sensory modalities (visual versus auditory) on brain dynamics and subjective experience, with visual stimulation reducing the measured relationship between neural entropic changes and subjective reports. Together, these findings support the choice of music — in contrast to visual stimulation — to modulate and support psychedelic therapy (19, 39, 40).

It remains possible that environments or stimuli different from the ones considered in this study could potentially lead to different results. Additionally, there are a number of phenomena relevant to the psychedelic experience for which having eyes open may be more conducive (e.g. feelings of communitas, or acute connection with nature (41)), which cannot be assessed within the current experimental design. Furthermore, the observed disruption between psychological phenomena and brain dynamics was only assessed via LZ applied to MEG data, and might not be true for other neural signatures.

Importantly, this study reveals that the effects of contextual elements on brain dynamics can be effectively tracked via current neuroimaging techniques. Our results establish LZ as a marker that is sensitive to the interaction between drug and context, which opens the door to future studies that may assess the effect of contextual elements on the brain during psychedelic therapy. This study, therefore, serves as a proof-of-concept translational investigation in healthy subjects, setting a precedent for future studies in clinical populations. Accompanying extensions into clinical populations, future work is also needed to further clarify how interactions between drug and context manifest on a psychological and neurobiological level, and how they can be harnessed for best therapeutic outcomes.

## Materials and Methods

### Data pre-processing

Data was collected with a 271-gradiometer CTF MEG scan. In addition, structural MRI scans of every subject were obtained for later inter-subject co-registration. Three subjects could not complete all stages of recording, or had excessive movement artefacts and were removed from the analysis altogether.

All pre-processing steps were performed using the FieldTrip toolbox (42). First, artefacts were removed by visual inspection and muscle and line noise effects were removed using ICA. Then we applied a 2^nd^-order lowpass Butterworth filter at 100 Hz and split the data into 2 s epochs for subsequent analysis.

For source reconstruction, we used the centroids of the AAL-90 atlas (43). The positions of these centroids were non-linearly inverse-warped to subject-specific grids using the subjects’ structural MRI scans, and source time series (a.k.a. *virtual sensors*) were estimated with a regularised LCMV beamformer. We calculated Lempel-Ziv complexity on these locations, and finally mapped them back onto the standard template for statistical analysis and visualisation. In addition, for the visualisation in Fig. 1c we computed LZ in sources reconstructed in a uniform 10 mm 3D grid.

### Lempel-Ziv complexity

The main tool of analysis used in this study is the Lempel-Ziv complexity (referred to as LZ), which estimates how diverse the patterns exhibited by a given signal are (44). The method was introduced by Abraham Lempel and Jacob Ziv to study the statistics of binary sequences (44), and was later extended (45, 46) to become the basis of the well-known “zip” compression algorithm. This algorithm has been used to study the diversity of patterns in EEG activity for more than 20 years, with some early studies focusing on epilepsy (47) and depth of anaesthesia (48).

LZ is calculated in two steps. First, the value of a given signal *X* of length *T* is binarised, calculating its mean value and turning each data point above it to “1”s and each point below it to “0”s. Then, the resulting binary sequence is scanned sequentially looking for distinct structures or “patterns.” Finally, the signal complexity is determined by the number of patterns found, denoted by *C*_LZ_(*X*). Regular signals can be characterized by a small number of patterns and hence have low *C*_LZ_, while irregular signals contain many different patterns and hence have a high *C*_LZ_.

Following the reasoning above, the LZ method identifies signal complexity with *richness of content* (49) — a signal is considered complex if it is not possible to provide a brief (i.e. compressed) representation of it. Accordingly, a popular way of understanding LZ is as a proxy for estimating the Kolmogorov complexity, the length of the shortest computer program that can reproduce a given pattern (50). However, we (and others) argue that this view is brittle in theory and of limited use in practice (51). A simpler and more direct interpretation of LZ is to focus on the quantity

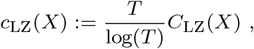

which is an efficient estimator of the entropy rate of *X* (52). The entropy rate measures how many bits of innovation are introduced by each new data sample (53), and is related with how hard it is to predict the next value of a sequence.^§^ This makes this normalised LZ, *c*_LZ_, a principled, data-efficient estimator of the diversity of the underlying neural process. For simplicity, the rest of the manuscript refers to *c*_LZ_ generically as LZ.

Unlike in previous studies, we do not apply a Hilbert transform, and instead apply the LZ procedure to the source-reconstructed, broadband signal. While there are certain interpretability advantages to using a Hilbert transform (for example, signal can be interpreted as the amplitude of an underlying neural oscillation), the Hilbert transform cannot be meaningfully applied to broadband signals, and pre-filtering the data would add further (undesired) degrees of freedom to our analysis. In practice, however, LZ is a remarkably robust measure and the same qualitative results hold under different pre-processing techniques. See Refs. (9, 15, 55) for further discussion.

In terms of algorithm, we follow the original procedure presented in Ref. (44) — commonly known as LZ76 — following the simplified algorithm described by Kaspar and Schuster (56). We note that although other versions of the LZ algorithm can also be employed to estimate the entropy rate (e.g. the common dictionary-based implementation (45, 46)), their computation time and convergence is slower than LZ76, making the latter a better choice for our experiments.

### Statistical modelling

To explore the effect of external conditions in detail, disentangling the effect of stimuli versus an effect *beyond* merely opening one’s eyes, the following encoding was used for the four conditions:

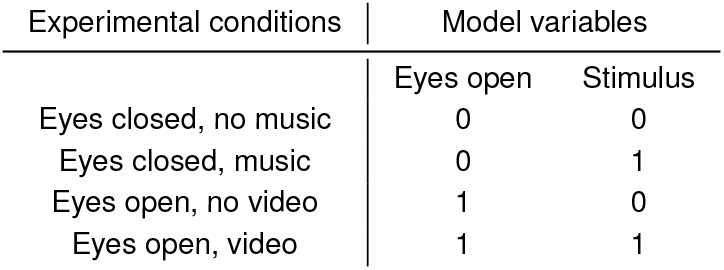

| Experimental conditions | Model variables |  |
| --- | --- | --- |
|  | Eyes open | Stimulus |
| Eyes closed, no music | 0 | 0 |
| Eyes closed, music | 0 | 1 |
| Eyes open, no video | 1 | 0 |
| Eyes open, video | 1 | 1 |

The paper considers various linear mixed-effects (LME) models, in most cases with a measure of interest (VAS ratings or LZ complexity) as target; drug, stimulus and eyes open as fixed effects; and subject identity as random effect. When constructing a model, all possible pairwise interactions were considered; then model selection is performed using the Bayesian Information Criterion (BIC). All the reported models correspond to the one selected by BIC. All models were estimated via restricted maximum likelihood, using the open-source packages `lme4 v.1.1-21` (57) and `lmerTest v.3.1-1` (58) on `R v.3.6.0`.

### Brain regions of interest

For the neural-psychometric correlation analysis in Fig. 3 onwards, we calculated the average LZ of several brain Regions of Interest (ROIs), each of them composed of a number of sub-regions represented in the AAL-90 atlas (43). For each subject, the mean LZ value of each ROI was obtained by averaging the LZ values of the source-reconstructed activity at the centroid of each sub-region.

For all the analyses, the following ROIs were considered: two sensory areas, two related to the DMN, one related to interoception and one to emotion. Specifically, the considered ROIs and their corresponding AAL-90 sub-regions are:

- Auditory: left and right Heschl areas;
- Visual: left and right Calcarine, bilateral Lingual, Cuneus, inferior, middle and supperior occipital;
- Amygdala: both left and right;
- Insula: both left and right;
- mPFC: left and right medial superior frontal gyrus; and
- Posterior DMN: bilateral posterior and median cingulate gyrus, middle temporal gyrus, and angular gyrus.

### Conditional predictive power analyses

The analyses of conditional predictive power were carried out according to the following procedure. Consider the case of studying the conditional predictive power of a given ROI *R*_1_ with respect to a particular subjective report *V*. We say that the predictive power from *R*_1_ to *V* is statistically mediated by another ROI *R*_2_ if (i) both *R*_1_ and *R*_2_ are significantly correlated with *V* according to their respective LME model (i.e. the FDR-corrected *p*-value of their estimated *β* is below 0.05); and (ii) when calculating a BIC-optimal LME model with *V* as target and both *R*_1_ and *R*_2_ as predictors, then the estimate of the effect of *R*_1_ loses significance (i.e. its *p*-value goes above 0.1, non-FDR-corrected). The diagrams in Figure 5a are built following this procedure in an iterative fashion, adding a link from *R*_1_ to *R*_2_ if and only if *R*_2_ mediates the predictive power of *R*_1_ about *V*, and a link from *R*_1_ to *V* if and only if it has unmediated predictive power about *V*. Figure 5b was obtained by an analogous procedure, where the roles of ROI and VAS were reversed.

### Ethics statement

This study was approved by the National Research Ethics Service committee London-West London and was conducted in accordance with the revised declaration of Helsinki (2000), the International Committee on Harmonization Good Clinical Practice guidelines, and National Health Service Research Governance Framework. Imperial College London sponsored the research, which was conducted under a Home Office license for research with schedule 1 drugs.

## ACKNOWLEDGMENTS

P.M. would like to thank Andrew Olson for discussion of the experimental results. P.M. and D.B. are funded by the Wellcome Trust (grant no. 210920/Z/18/Z). F.R. is supported by the Ad Astra Chandaria foundation. A.B. and A.S. are grateful to the Dr. Mortimer and Theresa Sackler Foundation, which supports the Sackler Centre for Consciousness Science. The initial study and data collection was supported by the Beckley Foundation as part of the Beckley-Imperial research programme, and by supporters of the Walacea.com crowdfunding campaign.

## Footnotes

* Entropy is understood here not as a thermodynamic but as an informational property, measuring the complexity of neural dynamics and the diversity of their configuration repertoire (see Methods).

† Although note that the role of auditory regions and insula is reversed for simple and complex imagery, respectively.

‡ The relation between LZ and spectral changes can be disentangled with more elaborate statistical methods (29), although this analysis is beyond the scope of this paper.

§ In effect, the mean entropy rate divided by two approximates the probability of making an error with the best informed guess about the next sample (54).

## Notes

The authors declare no conflict of interest.

### Competing Interest Statement

The authors have declared no competing interest.

