## Supplementary Material for "Effects of external stimulation on psychedelic state neurodynamics"

#### This PDF file includes:

Supplementary text
Figs. S1 to S5
Tables S1 to S10
References for SI reference citations

### Supporting Information Text

### Additional controls on the interaction between drug and condition

In Figure 1 of the main text, we report as the main finding of the paper the result that there exists a negative drug×eyes interaction effect on whole-brain average LZ. To confirm these results we performed a number of additional controls (all agreeing with our conclusions), described here and reported in the tables below:

(1) Table S1 presents the results of a LME model in which the condition variable is an integer coding of external stimulation, according to the following order: closed (0), music (1), open (2), and video (3). For the LME, the condition was treated as a continuous variable. This is the model shown in Figure 1b of the main text.

(2) Table S2 shows the result of a two-way ANOVA run on the drug and condition variables, with the condition variable being a four-valued discrete variable.

(3) Table S3 presents the replication of the results in Table 1 of the main text on data under a more conservative filtering — a low-pass $4^{\text{th}}$-order Butterworth filter with cut-off frequency at 30 Hz.

(4) Table S4 presents the results of a LME model controlling for ordering effects between the stimulus and non-stimulus sessions — i.e. run on a different set of data in which the eyes-closed and eyes-open conditions were recorded after the music and video conditions, respectively.

### Sensitivity of correlations to the external condition

Here we provide additional information about the results in Figure 3 of the main text, related to how external stimulation affects correlation (i) between VAS scores and LZ in the considered ROIs, and (ii) between LZ in each pair of ROIs. To study this, we computed the between-subjects correlation between VAS and LZ changes (LSD - placebo), in each experimental condition (Figure S1). To draw quantitative conclusions, we constructed a multivariate regression model using the correlation coefficients as target variables and stimuli and eye opening as predictors. We constructed two separate models: one using VAS-VAS correlations as target (Table S5), and one using VAS-ROI correlations as target (Table S6).

In order to discard potential ordering effects driving this results, we re-ran these analyses on the same order-flipped data used in item (4) above, with nearly identical results (Figure S2). The numerical results of these models are presented in Tables S7 and S8.

### Relation between LZ complexity and alpha power

Decreased power in the alpha band is one of the most systematic spectral effects reported in the M/EEG psychedelics literature (1–3). This has raised questions regarding to what extent is the LSD-induced increase in LZ merely a reflection of the corresponding alpha power suppression. To touch briefly on this issue, here we show that while alpha suppression shows a behaviour similar to LZ when comparing across gross state changes (Figures S3 and S4), it is substantially less correlated with subjective reports than LZ (Figure S5).

More specifically, first Figure S3 presents a comparison between LZ and alpha power for each subject in the placebo session, for each of the four experimental conditions. As expected, alpha power is inversely related to LZ, with richer stimuli inducing stronger suppression. Additionally, Figure S4 shows that when measured with alpha suppression the drug also has an interaction with external stimulation (c.f. Figure 1b in the main text). Quantitatively, LME modelling reveals that both the drug and eye opening have significant negative effects on alpha power, and that they have a significant positive interaction (Table S10).

Finally, we explored the preditive capability of alpha changes in all ROIs to predict VAS ratings (Figure S5). In this case, unlike the above, the results of alpha suppression do not resemble those of LZ: while some relationships survive the multiple-comparisons correction, alpha power seems to be far less capable of predicting subjective reports than LZ (c.f. Figure 4 in the main text). This suggests LZ to be a more sensitive, and therefore more suitable, marker of the psychedelic experience — although a full analysis decomposing the spectral components of the LZ difference is needed to shed more light on the matter (4).

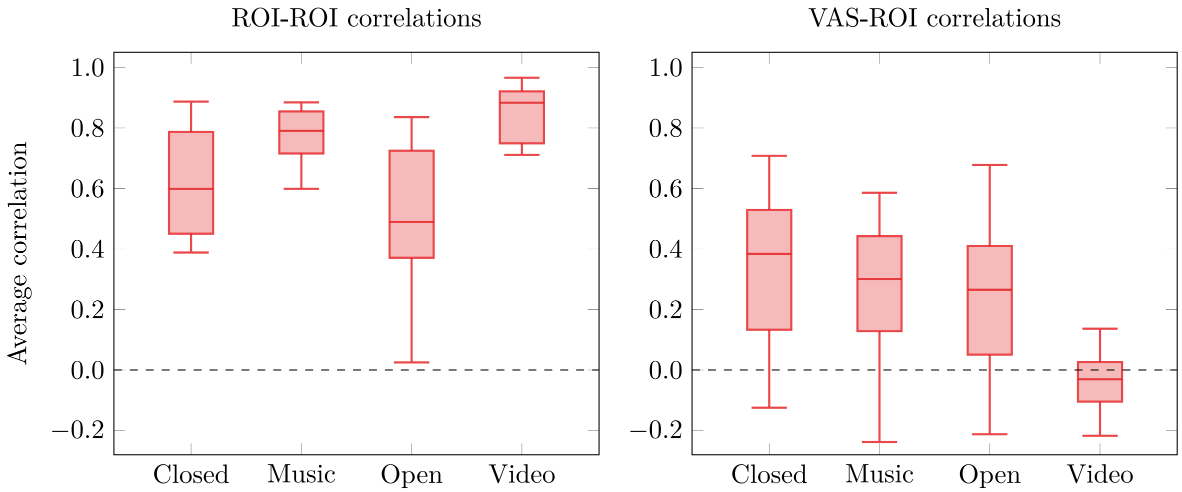


**Fig. S1.** Between-subjects correlation between pairs of ROIs (left) and ROIs and VAS items (right).

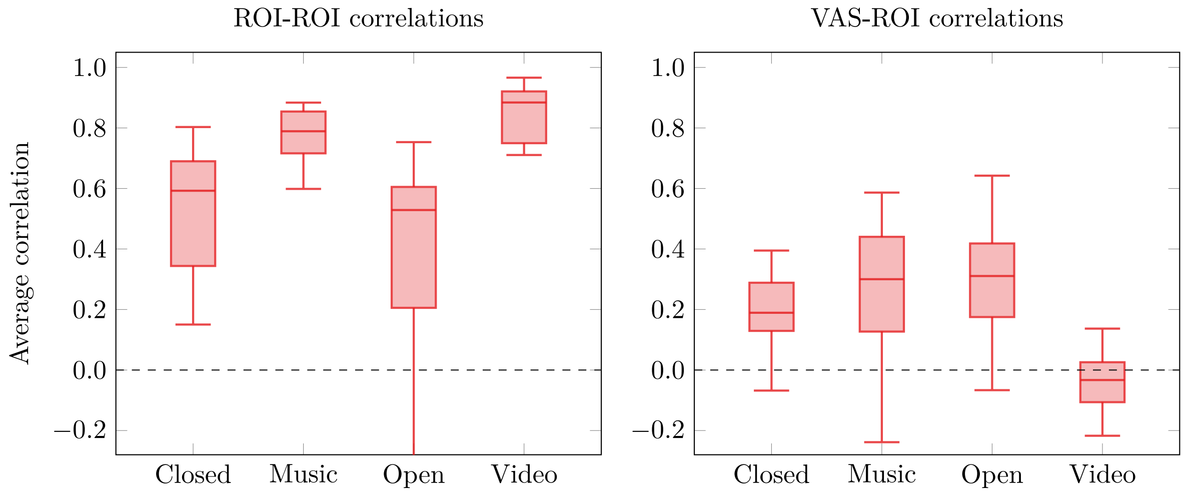


**Fig. S2.** Between-subjects correlation between pairs of ROIs (left) and ROIs and VAS items (right), controlling for ordering effects.

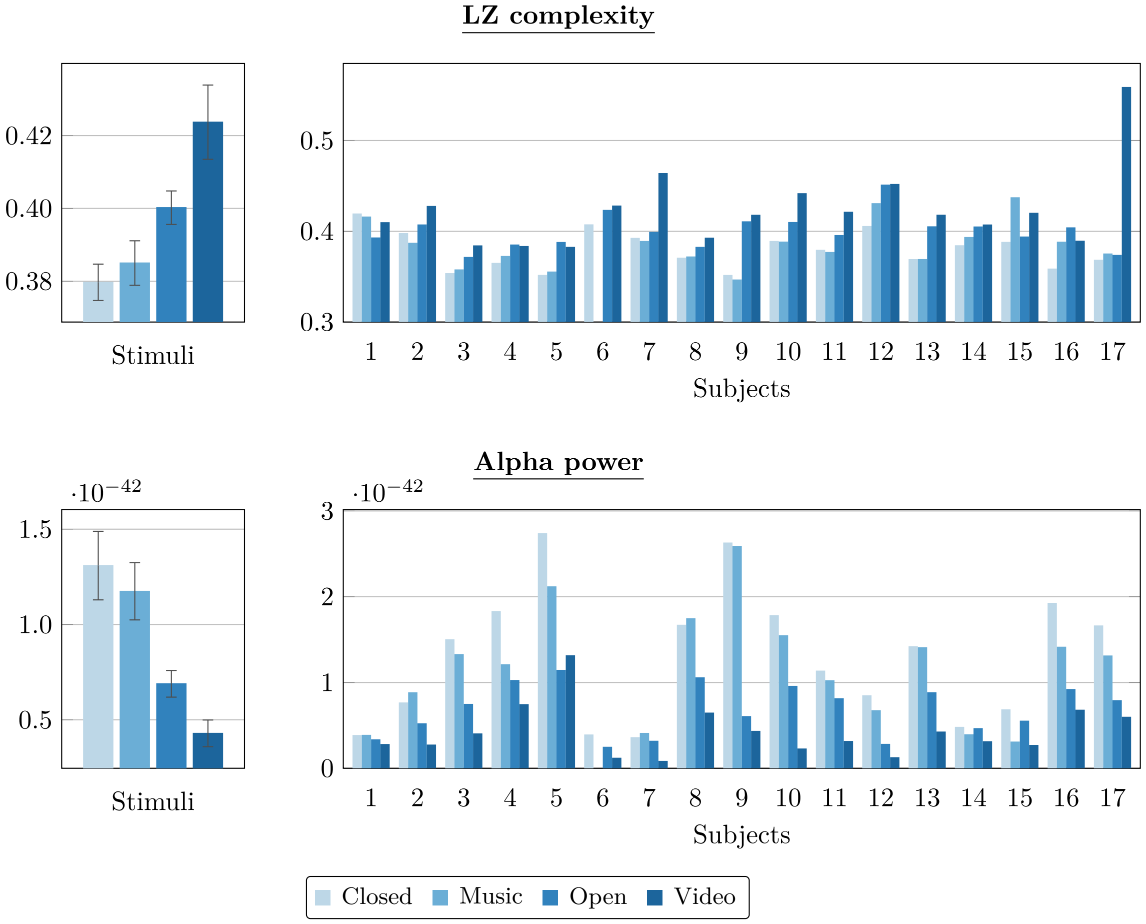


**Fig. S3.** Data from the non-drug placebo condition: External stimulation increases LZ complexity (top) and supresses alpha power (bottom). For each measure, data are provided per subject (right) and averaged across subjects (left; error bar is standard error). LZ complexity is reported in bits/sample, and alpha power in $T^2$.

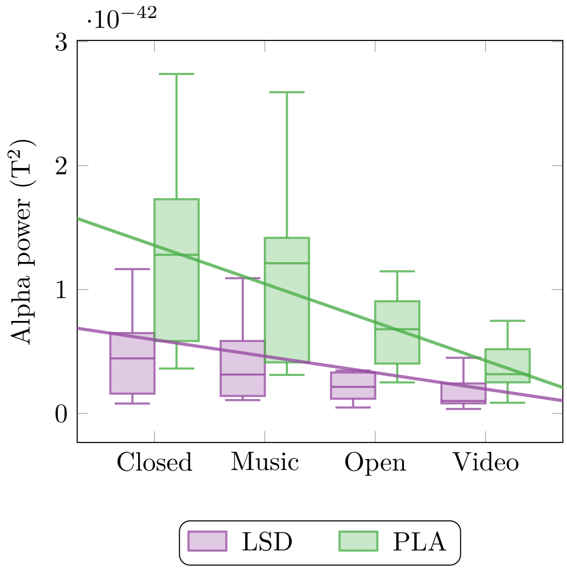


**Fig. S4.** Whole-brain average alpha (8-13 Hz) power for both LSD and placebo sessions, for all experimental conditions.

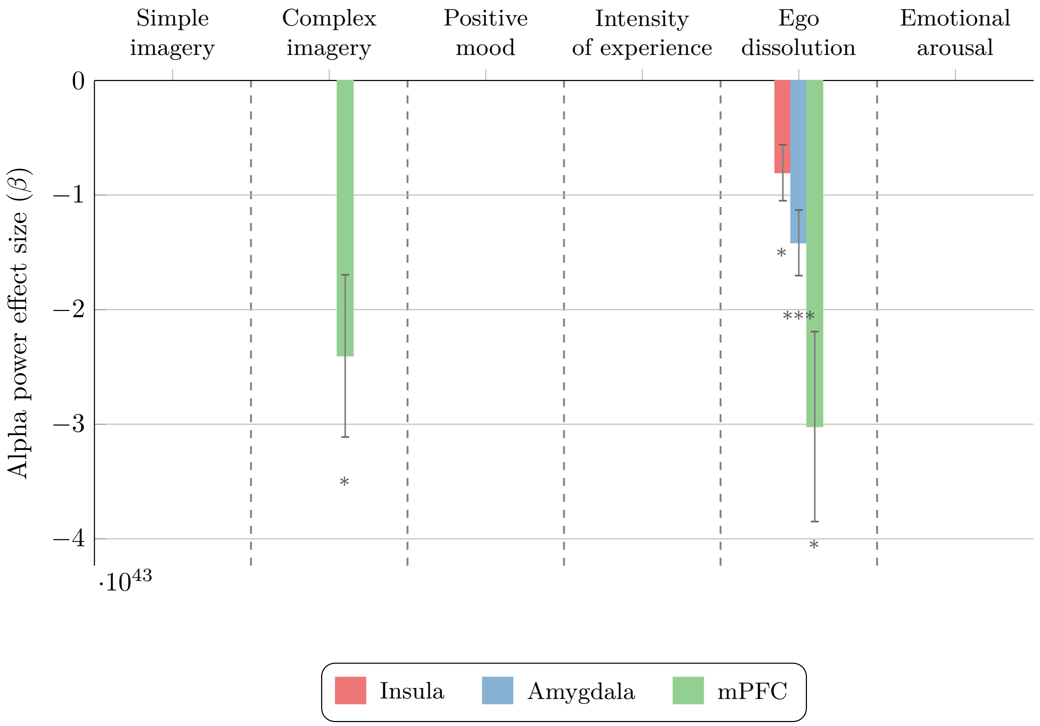


**Fig. S5.** Estimates, standard error, and FDR-corrected statistical significance ($^{*}$: $p < 0.05$,$^{**}$: $p < 0.01$,$^{***}$: $p < 0.001$), of the effect of the alpha power differences (LSD-PLA), averaged over various ROIs, for predicting VAS differences (LSD-PLA). ROIs not shown had no significant effect from alpha power on any VAS item.

**Table S1. Coefficients of LME model predicting average LZ with an integer-coded condition variable.**

| | $\beta$ | SE | $p$ |
|---|---|---|---|
| **Drug** | 0.050 | 0.005 | <0.001 |
| **Condition** | 0.012 | 0.002 | <0.001 |
| **Drug×Condition** | -0.007 | 0.002 | 0.009 |

**Table S2. Results of a two-way ANOVA on average LZ using an integer-coded condition variable.**

| | Mean Square | F | $p$ |
|---|---|---|---|
| **Drug** | 0.050 | 107.4 | <0.001 |
| **Condition** | 0.013 | 27.0 | <0.001 |
| **Drug$\times$Condition** | 0.002 | 5.0 | 0.027 |

**Table S3. Coefficients of LME model predicting average LZ on data low-pass filtered at 30 Hz.**

| | $\beta$ | SE | $p$ |
|---|---|---|---|
| **Drug** | $8.44 \times 10^{-3}$ | $9.67 \times 10^{-4}$ | <0.001 |
| **Stimulus** | $1.60 \times 10^{-3}$ | $6.80 \times 10^{-4}$ | 0.021 |
| **Eyes** | $5.28 \times 10^{-3}$ | $9.61 \times 10^{-4}$ | <0.001 |
| **Drug×Eyes** | $-4.56 \times 10^{-3}$ | $1.36 \times 10^{-3}$ | 0.001 |

**Table S4. Coefficients of LME model predicting average LZ controlling for ordering effects.**

| | $\beta$ | SE | $p$ |
|---|---|---|---|
| **Drug** | 0.044 | 0.004 | <0.001 |
| **Stimulus** | 0.009 | 0.003 | 0.005 |
| **Eyes** | 0.021 | 0.004 | <0.001 |
| **Drug$\times$Eyes** | -0.012 | 0.006 | 0.048 |

**Table S5. Coefficients of multivariate linear model for predicting VAS-ROI correlation. Adjusted-R=0.28, F(3,116)=16.59, p<0.001.**

| | $\beta$ | SE | $p$ |
|---|---|---|---|
| **Stimulus** | - 0.07 | 0.05 | 0.168 |
| **Eyes** | - 0.08 | 0.05 | 0.127 |
| **Stimulus$\times$Eyes** | -0.21 | 0.08 | 0.006 |

**Table S6. Coefficients of multivariate linear model for predicting ROI-ROI correlation. Adjusted-R=0.42, F(3,116)=29.89, p<0.001.**

| | $\beta$ | SE | $p$ |
|---|---|---|---|
| **Stimulus** | 0.15 | 0.04 | <0.001 |
| **Eyes** | -0.10 | 0.04 | 0.011 |
| **Stimulus×Eyes** | 0.18 | 0.06 | 0.001 |

**Table S7. Coefficients of multivariate linear model for predicting VAS-ROI correlation controlling for ordering effects. Adjusted-R=0.27, F(3,116)=15.63, p<0.001.**

| | $\beta$ | SE | $p$ |
|---|---|---|---|
| **Stimulus** | - 0.07 | 0.04 | 0.085 |
| **Eyes** | - 0.07 | 0.05 | 0.139 |
| **Stimulus$\times$Eyes** | -0.37 | 0.07 | <0.001 |

**Table S8. Coefficients of multivariate linear model for predicting ROI-ROI correlation controlling for ordering effects. Adjusted-R=0.47, F(3,116)=35.91, p<0.001.**

| | $\beta$ | SE | $p$ |
|---|---|---|---|
| **Stimulus** | 0.24 | 0.05 | <0.001 |
| **Eyes** | -0.15 | 0.05 | 0.004 |
| **Stimulus$\times$Eyes** | 0.23 | 0.07 | 0.002 |

**Table S9. Model FDR-corrected $p$-value and coefficients (mean $\pm$ SE) of LME models predicting VAS changes from LZ changes in each ROI. Models for all VAS,ROI pairs were fitted following the procedure in Materials and Methods, and those with $p < 0.05$ are shown here.**

| VAS | ROI | $p$-value | LZ | Stimulus | Eyes | LZ $\times$ Stimulus | LZ $\times$ Eyes |
|---|---|---|---|---|---|---|---|
| Simple imagery | Whole brain | $0.002$ | $116.4 \pm 34.9$ | $1.94 \pm 0.78$ | *n/a* | *n.s.* | *n/a* |
| | pDMN | $0.003$ | $111.6 \pm 31.9$ | $1.68 \pm 0.83$ | *n/a* | *n.s.* | *n/a* |
| | Insula | $< 0.001$ | $85.8 \pm 20.1$ | $42.51 \pm 0.68$ | *n/a* | *n.s.* | *n/a* |
| | Visual | $< 0.001$ | $135.1 \pm 31.2$ | $1.94 \pm 0.77$ | *n/a* | *n.s.* | *n/a* |
| | Auditory | $0.032$ | $63.4 \pm 26.0$ | $1.45 \pm 0.89$ | *n/a* | *n.s.* | *n/a* |
| Complex imagery | Whole brain | $0.012$ | $94.0 \pm 34.2$ | $0.02 \pm 1.04$ | *n/a* | *n.s.* | *n/a* |
| | pDMN | $0.006$ | $94.9 \pm 28.7$ | $-0.20 \pm 1.01$ | *n/a* | *n.s.* | *n/a* |
| | Insula | $0.023$ | $55.7 \pm 21.3$ | $0.36 \pm 1.04$ | *n/a* | *n.s.* | *n/a* |
| | Visual | $0.006$ | $109.2 \pm 30.9$ | $-0.002 \pm 0.89$ | *n/a* | *n.s.* | *n/a* |
| | Auditory | $0.006$ | $77.1 \pm 21.4$ | $-0.51 \pm 1.05$ | *n/a* | *n.s.* | *n/a* |
| Positive mood | Whole brain | $0.027$ | $50.6 \pm 22.2$ | $1.19 \pm 0.91$ | $-1.70 \pm 0.96$ | *n.s.* | *n.s.* |
| | Amygdala | $0.020$ | $44.5 \pm 15.9$ | $1.15 \pm 0.89$ | $-1.62 \pm 0.94$ | *n.s.* | *n.s.* |
| | Visual | $0.020$ | $58.0 \pm 19.4$ | $1.35 \pm 0.90$ | $-1.85 \pm 0.91$ | *n.s.* | *n.s.* |
| Intensity | Visual | $0.047$ | $66.7 \pm 23.6$ | $1.06 \pm 0.67$ | $2.54 \pm 1.24$ | *n.s.* | $-81.1 \pm 25.1$ |
| Ego dissolution | Whole brain | $0.010$ | $115.7 \pm 36.3$ | $3.74 \pm 1.66$ | $-0.38 \pm 0.84$ | $-110.1 \pm 37.1$ | *n.s.* |
| | pDMN | $0.005$ | $114.01 \pm 30.0$ | $3.67 \pm 1.47$ | $-0.65 \pm 0.79$ | $-107.9 \pm 31.4$ | *n.s.* |
| | Insula | $0.018$ | $59.9 \pm 18.2$ | $2.16 \pm 1.40$ | $-0.24 \pm 0.84$ | $-53.6 \pm 21.8$ | *n.s.* |
| | Auditory | $0.003$ | $88.8 \pm 21.2$ | $4.07 \pm 1.44$ | $-0.78 \pm 0.78$ | $-87.7 \pm 23.1$ | *n.s.* |

**Table S10. Coefficients of LME model predicting average Alpha power change.**

| | $\beta$ | SE | $p$ |
|---|---|---|---|
| **Drug** | $-6.31 \times 10^{-43}$ | $6.81 \times 10^{-44}$ | $< 0.001$ |
| **Stimulus** | $-1.40 \times 10^{-43}$ | $4.79 \times 10^{-44}$ | $0.004$ |
| **Eyes** | $-6.17 \times 10^{-43}$ | $6.77 \times 10^{-44}$ | $< 0.001$ |
| **Drug×Eyes** | $3.57 \times 10^{-43}$ | $9.56 \times 10^{-44}$ | $< 0.001$ |
